# From rare Copy Number Variations to biological processes in ADHD

**DOI:** 10.1101/762419

**Authors:** Benjamin Harich, Monique van der Voet, Marieke Klein, Pavel Čížek, Michaela Fenckova, Annette Schenck, Barbara Franke

## Abstract

**Aim:** Attention-deficit/hyperactivity disorder (ADHD) is a highly heritable psychiatric disorder. The objective of this study was to define ADHD-associated candidate genes, and their associated molecular modules and biological themes, based on the analysis of rare genetic variants.

**Methods:** We combined data from 11 published copy number variation (CNV) studies in 6176 individuals with ADHD and 25026 controls and prioritized genes by applying an integrative strategy based on criteria including recurrence in ADHD individuals, absence in controls, complete coverage in copy number gains, and presence in the minimal region common to overlapping CNVs, as well as on protein-protein interactions and information from cross-species genotype-phenotype annotation.

**Results:** We localized 2241 eligible genes in the 1532 reported CNVs, of which we classified 432 as high-priority ADHD candidate genes. The high-priority ADHD candidate genes were significantly co-expressed in the brain. A network of 66 genes was supported by ADHD-relevant phenotypes in the cross-species database. In addition, four significantly interconnected protein modules were found among the high-priority ADHD genes. A total of 26 genes were observed across all applied bioinformatic methods. Look-up in the latest genome-wide association study for ADHD showed that among those 26, *POLR3C* and *RBFOX1* were also supported by common genetic variants.

**Conclusions:** Integration of a stringent filtering procedure in CNV studies with suitable bioinformatics approaches can identify ADHD candidate genes at increased levels of credibility. Our pipeline provides additional insight in the molecular mechanisms underlying ADHD and allows prioritization of genes for functional validation in validated model organisms.

## Introduction

Attention-deficit/hyperactivity disorder (ADHD) is one of the most common neuropsychiatric disorders, with a prevalence of 5–6% in children.(1) The disorder persists into adulthood in a significant proportion of affected individuals, resulting in a prevalence of 2.5–4.9% in adults.(2) The clinical symptoms of ADHD include age-inappropriate inattention, hyperactivity, and impulsivity.(3) Twin and adoption studies estimated a high heritability of 76% for ADHD.(3)

Identification of the genes implicated in ADHD and their molecular functions offers opportunities to understand the neurobiological mechanisms leading to ADHD and facilitates the development of diagnostic tools and new treatments. However, despite the high heritability, identification of ADHD risk genes has been difficult, mainly due to ADHD’s complex genetic architecture.(2,4,5) To date, mainly genetic variants that frequently occur in the population have been investigated for their role in ADHD, either through studies of candidate genes or hypothesis-free genome-wide association studies (GWAS).(6,7) A recent GWAS meta-analysis identified the first 12 loci harbouring ADHD risk variants.(8) Another type of GWAS has focused on the association of rare copy number variants (CNVs) with ADHD. Such CNV GWASs have largely concentrated on rare events (primarily) observed in individuals diagnosed with ADHD. We analysed the 11 studies published to date that have detected rare CNVs in ADHD cases.(9,10,19,11–18) Those CNV GWASs implicated more than 2200 candidate genes in ADHD, though most have investigated rather limited sample sizes, and most of the CNVs were detected only in single patients. Based on the average mutation rate in-between individuals in the general population, single-patient rare findings have a high chance to be false positive; it is thus important to concentrate on repeatedly occurring copy number events.(20)

Data integration from various sources is an important strategy to move from genes to biologically meaningful modules. Examples for this come from publications on ADHD and related disorders.(21),(22) A publication on autism spectrum disorders showed how data integration enabled the identification of highly conserved gene clusters that improve our understanding of neuropsychiatric disorders.(21) Similarly, a recent study found a significant overlap of ADHD case CNVs with targets of the Fragile-X mental retardation protein (FMRP), a gene cluster involved in neurodevelopmental disorder risk.(22) In many cases, data integration currently takes place only across data modalities derived from studies in humans. This neglects the wealth of phenotypic information that can be derived from model organisms such as monkey, rat, mouse, zebrafish, and fruit fly.(7)

In this study, we surveyed and integrated data on CNVs associated with ADHD from existing publications, aiming to define robustly ADHD-associated genes, molecular modules, and biological themes underlying this disorder. We combined data from the 11 published ADHD CNV studies and applied an integrative strategy using redundancy criteria, data on protein-protein interactions, and – most innovative – employing information from cross-species genotype-phenotype annotation to prioritize candidate genes.(23,24) We classified 432 high-priority ADHD candidate genes, supported by co-expression, cross-species phenotype, and protein interaction information, with 26 genes highlighted across all approaches. Integration with data on common genetic variants showed that, amongst these 26 genes, *POLR3C* and *RBFOX1* were significantly associated with ADHD in the largest SNP-based GWAS meta-analysis to date.

## Materials and Methods

### Identification of genes affected by ADHD CNVs

Coordinates of rare CNVs occurring in individuals diagnosed with ADHD (cases) were retrieved from 11 studies published until now (9,10,19,11–18) Discovery samples of in total 6176 ADHD cases and 25026 controls provided the bases for the analysed studies (see **Supplementary Table 1** for study characteristics). Coordinates of 1532 CNVs were retrieved for our analysis. These were mapped to the same reference human genome (hg19) using UCSC Lift Genome Annotations; the minimal ratio of bases that must remap was set to 0.95.(25) The CNV coordinates were used to retrieve RefGene information from the UCSC MySQL database, using a Structured Query Language (SQL) query (see **Supplementary Text 1**).(26) Information retrieved included the overlap with coding sequence, transcriptional direction, the total gene and CNV size in base pairs, exact gene start/end position, and percentage of gene coding sequence (CDS) represented by the CNV (**Supplementary Table 1**, Tab 1).

### Selection of genes recurrently affected in ADHD CNVs

Transcript variants and biotypes were extracted for each CNV through a batch NCBI nucleotide query (**Supplementary Table 2**, Tab 1 for biotypes). For gene copy-number losses, we included all those genes entirely deleted or partially truncated. We also annotated, whether the N- or C-terminal region of transcripts were affected (**Supplementary Table 2**, Tab 1). For gene copy-number gains, we considered transcripts for which both a 2kb promoter and the coding region were entirely duplicated. For overlapping CNVs, the minimal region of overlap was identified to narrow down the putative region involved in ADHD. Only mRNA-coding genes affected by CNVs in at least two cases with ADHD were selected for subsequent analyses (high-priority catalogue), given interpretability and possibility to perform cluster and protein-protein interaction analyses.

### Co-expression network analysis

The BrainSpan developmental transcriptome data set (RNA-Seq Gencode v10) was used to investigate the overrepresentation of co-expressed genes of our high priority gene-set across all brain regions and developmental time points (embryo to adult), relative to the rest of the genome.(27) The expression coefficients for each mRNA-coding gene in all time points and brain regions in the BrainSpan data set were concatenated. The co-expression correlation score was calculated for each gene-pair. Gene-pairs with correlation score >0.3 were assigned to a co-expression network, each node representing a single gene and each connection representing the correlation score. The sum of the correlation scores for the investigated gene-set and for 10000 random gene-sets of the same size was calculated. An enrichment score was calculated by dividing the sum of the correlation scores per gene-set by the mean of the 10000 random gene-sets. P-values were calculated by comparing how many of the 10000 correlation scores of the random gene-sets were equal or higher than those of the investigated gene-set.

### Integrated cross-species phenotype and protein-protein interaction network

We used the Monarch Initiative cross-species phenotype database to retrieve genes associated with an ADHD-related phenotype.(28) We defined core phenotypes, by selecting terms based on attention-deficit, hyperactivity, and impulsivity (**Supplementary Table 3**). The genes connected to the cross-species core phenotypes of ADHD were subsequently superimposed to inter-species BioGRID protein interaction data. The interaction plot was visualised with Cytoscape 3.4.0.(29)

### Identification of enriched-protein interaction modules

Networks of physical interactions in the gene-set were assessed with DAPPLE 0.17(30,31) based on InWeb data with the following parameter settings: Number of permutations: 1000 with adaptive permutation function, Plot: true, Seed File: genes. A set of 432 high-priority genes formed the input for this analysis. The modules were visualised with Cytoscape 3.4.0.(29)

### Gene-based and gene-set association analyses of prioritized modules with ADHD risk

We used data from the recent meta-analysis of genome-wide association studies (GWAS) of 20,183 patients with ADHD and 35,191 controls as performed by the Psychiatric Genomics Consortium (PGC) ADHD Working Group and the Danish iPSYCH Initiative.(8) Details on the samples and quality control can be found in the **Supplementary Methods** and in Demontis et al..(8)

Gene-based association analyses were performed using the Multi-marker Analysis of GenoMic Annotation (MAGMA) software package (version 1.05; for details see **Supplementary Text 1**).(32) Genes were considered gene-wide significant if they reached the Bonferroni correction threshold-adjusted for the number of genes tested (*p*<0.05/26).

## Results

### Identification of genes located in ADHD-associated CNVs and definition of high-priority gene-set

We extracted data from the 11 studies reporting rare CNVs in a total discovery sample of 6176 ADHD cases and 25026 controls (see **Supplementary Table 1** for study characteristics). Coordinates of 1532 CNVs were retrieved, containing 2241 mRNA-coding candidate genes.

To identify the genes with an increased likelihood of contributing to ADHD pathology, we removed all genes duplicated with an incomplete promoter or coding sequence, or those aberrations found in controls across all studies (Figure 1). Genes identified in at least two rare CNVs were placed among the high-ranking candidates, due to their recurring nature. In addition, we calculated the minimal region common to overlapping CNVs, to narrow down the region of interest. Together, this resulted in a high-ranking list of 432 genes (**Supplementary Table 2, High-priority gene list**). The remaining 1316 genes, observed in only a single patient, were considered low-ranking (**Supplementary Table 2, Low-priority gene list**).

**Figure 1.**
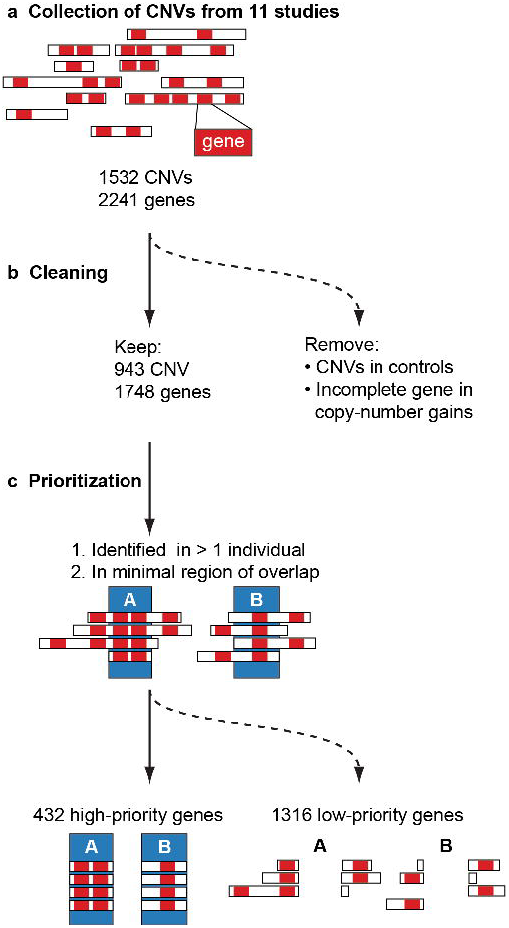
CNV processing scheme to establish high- and low-priority ADHD candidate genes. **(a)** Collection: CNV coordinates were extracted from 11 different studies reporting rare CNVs in ADHD cohorts (see **Supplementary Table 1**). **(b)** Cleaning: CNVs identified in controls and copy-number gains with an incomplete promoter and coding sequences were excluded. **(c)** Prioritisation: a high-priority gene list was generated by excluding genes found only in a single individual with ADHD and genes not present in the minimal overlapping region.

### High-priority ADHD candidate genes show increased co-expression in the brain

It has been shown that proteins encoded by genes implicated in a genetically heterogeneous disorder tend to operate in common molecular pathways and processes.(22,33–36) To evaluate biological coherence of high-priority ADHD candidate genes in an unbiased way, we assessed their co-expression, a prerequisite for genes to jointly act in biological and developmental processes, during the development of the most relevant tissue, the brain. We used the BrainSpan data set to test for gene co-expression and found a significant enrichment (E) of co-expressed genes in the high-priority gene-list (*n*=432; *E*=1.04, *p*=0.0044). The low-priority genes (*n*=1316; *E*=1.01, *p*=0.28) did not show significant co-expression enrichment.

### Cross-species phenotypes link a network of 66 high-priority candidate genes to ADHD core symptoms

We used the Monarch Initiative cross-species genotype–phenotype database, which contains phenotypic information from 58 species, to i) evaluate our gene prioritisation, ii) retrieve independent evidence for the relevance of our high-priority candidate gene-set for core ADHD features, and iii) identify functionally associated networks of high-priority genes with the ADHD core symptoms hyperactivity, attention-deficit, and impulsivity (for exact search terms see **Supplementary Table 3**).(28) Eighteen of the 432 high-priority ADHD CNV candidate genes were associated with cross-species terms related to attention and hyperactivity: attention-deficit/hyperactivity disorder, hyperactivity, increased vertical activity, hyperactive, and abnormally increased process quality locomotory exploration behaviour (Figure 2).

**Figure 2.**
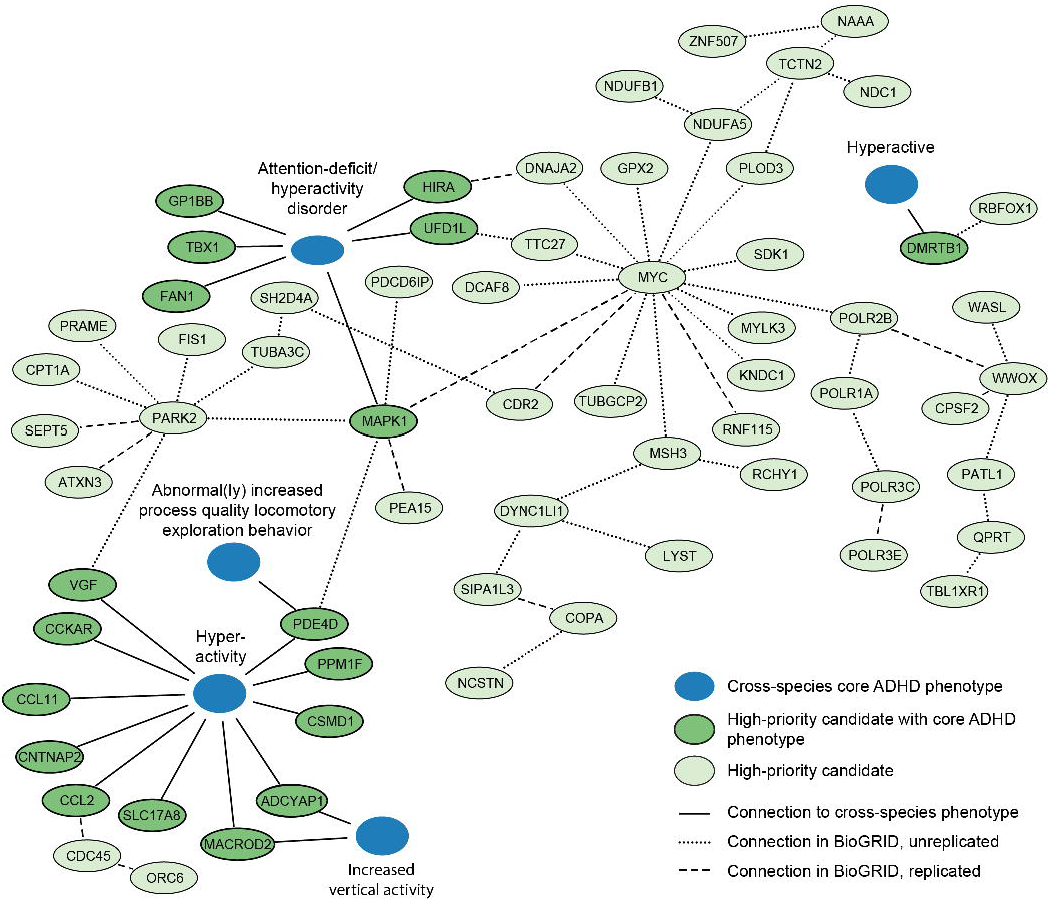
Cross-species phenotypes link an interaction network of 66 high-priority candidate genes to ADHD core symptoms. Of the 432 high-priority ADHD CNV candidate genes, 18 were found in the Monarch Initiative database to have a core ADHD phenotype relating to hyperactivity, attention deficit, and impulsivity. These 18 genes were used as seeds to create a BioGRID interaction network, using cross-species information on direct protein–protein interactions, genetic interactions and predicted interactions. We found a highly interconnected network with 48 secondary interactors from the high-priority gene list. Dashed lines show whether a connection was found once (^..^) or multiple times (--) in the BioGRID database.

Based on the 18 genes validated by the cross-species approaches, we mined the cross-species Biological General Repository for Interaction Datasets (BioGRID) for interactors. This approach connected 48 additional genes of our high-priority catalogue to the cross-species terms (Figure 2).

### CNV-derived high-priority ADHD candidate genes form molecular modules implicating specific biological processes in ADHD pathology

In addition to the cross-species approach, we also used the DAPPLE algorithm to identify significantly connected proteins among the 432 high-priority genes, based on the integration of human data from protein–protein, genetic, and pathway interactions.(30) This algorithm depicted 17 modules of connected proteins, each comprising of 2–15 ADHD high-priority candidates that directly interacted with each other (Figure 3). Taking both direct and indirect interactions into account, the hubs with significant connectivity contained 20 proteins (Figure 3 and **Supplementary Table 4**). Of those, eight significant proteins were found in four direct protein-interaction modules (Figure 3, **Module 1–4**). Of those modules, ten proteins, WWOX, PPM1F, PARK2, TUBA3C, MAPK1, MYC, SEPT5, POLR2B, POLR1A, and *POLR3C* were found connected to cross-species core ADHD phenotypes (Figure 2).

**Figure 3.**
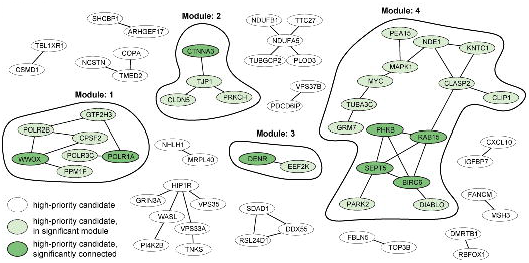
DAPPLE interaction network identifies four molecular modules with significantly connected proteins. A DAPPLE analysis was performed to identify highly related protein–protein interaction clusters based on direct- and indirect connectivity. Four modules containing significantly connected proteins were identified.

### Two genes were found significantly associated with ADHD in common variant GWA studies

A set of 26 genes amongst the original 432 high-priority CNV-derived ADHD candidate genes was consistently observed in all the different approaches we employed (Figure 4, Table 1). We performed gene-based association analyses for all 26 genes to evaluate whether these are also implicated in ADHD risk through common genetic variants, using the largest meta-analytic GWAS data for ADHD currently available (*n*=55374, PGC-iPSYCH ADHD working groups). Two genes were significantly associated with ADHD after correction for multiple testing. These genes were *POLR3C* (*p*= 0.000020373) and *RBFOX1* (*p*= 0.00018202) (see **Supplementary Table 5 and Supplementary Figure 1**). Interestingly, the *POLR3C* gene was among the top 0.43% of the most strongly associated genes (rank 80 out of 18411).

**Figure 4.**
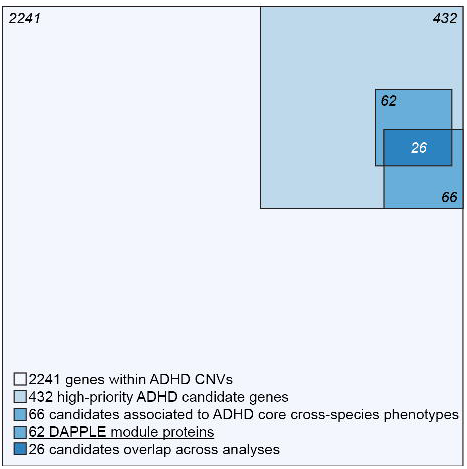
Visual representation of the gene-selection process. A summary of the analysis process undertaken in the paper, starting from 2241 genes prioritised to 432 high-priority candidates (Figure 1), followed by a cross-species phenotype and BioGRID analysis of 66 genes (Figure 2) and DAPPLE analysis with 62 candidates (Figure 3). In total 26 candidates overlap across analyses (Table 1).

**Table 1.**
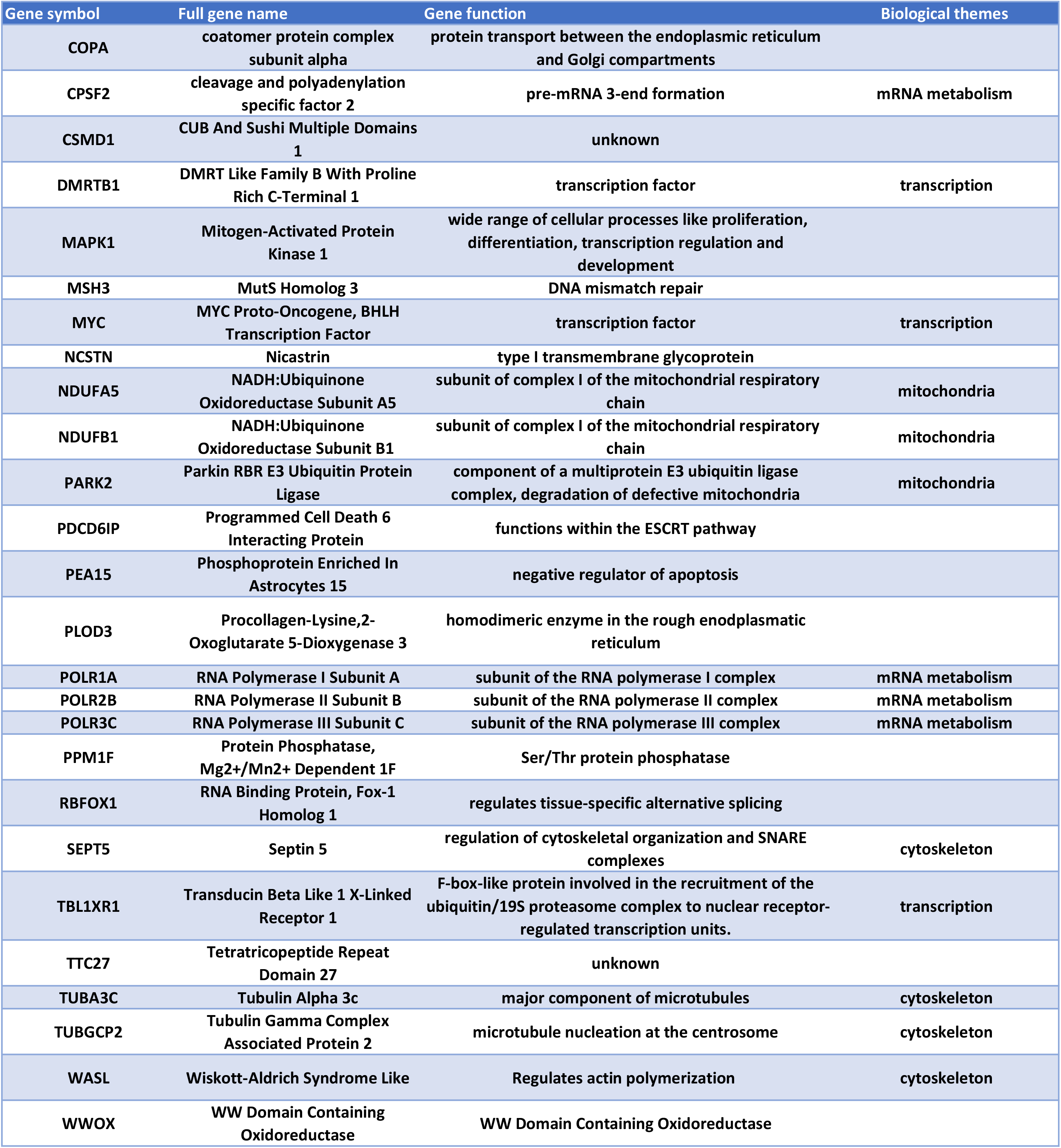
The 26 genes consistently identified across all bioinformatics approaches employed in the current study.

| Gene symbol | Full gene name | Gene function | Biological themes |
| --- | --- | --- | --- |
| COPA | coatomer protein complex subunit alpha | protein transport between the endoplasmic reticulum and Golgi compartments |  |
| CPSF2 | cleavage and polyadenylation specific factor 2 | pre-mRNA 3-end formation | mRNA metabolism |
| CSMD1 | CUB And Sushi Multiple Domains 1 | unknown |  |
| DMRTB1 | DMRT Like Family B With Proline Rich C-Terminal 1 | transcription factor | transcription |
| MAPK1 | Mitogen-Activated Protein Kinase 1 | wide range of cellular processes like proliferation, differentiation, transcription regulation and development |  |
| MSH3 | MutS Homolog 3 | DNA mismatch repair |  |
| MYC | MYC Proto-Oncogene, BHLH Transcription Factor | transcription factor | transcription |
| NCSTN | Nicastrin | type I transmembrane glycoprotein |  |
| NDUFA5 | NADH:Ubiquinone Oxidoreductase Subunit A5 | subunit of complex I of the mitochondrial respiratory chain | mitochondria |
| NDUFB1 | NADH:Ubiquinone Oxidoreductase Subunit B1 | subunit of complex I of the mitochondrial respiratory chain | mitochondria |
| PARK2 | Parkin RBR E3 Ubiquitin Protein Ligase | component of a multiprotein E3 ubiquitin ligase complex, degradation of defective mitochondria | mitochondria |
| PDCD6IP | Programmed Cell Death 6 Interacting Protein | functions within the ESCRT pathway |  |
| PEA15 | Phosphoprotein Enriched In Astrocytes 15 | negative regulator of apoptosis |  |
| PLOD3 | Procollagen-Lysine,2-Oxoglutarate 5-Dioxygenase 3 | homodimeric enzyme in the rough endoplasmic reticulum |  |
| POLR1A | RNA Polymerase I Subunit A | subunit of the RNA polymerase I complex | mRNA metabolism |
| POLR2B | RNA Polymerase II Subunit B | subunit of the RNA polymerase II complex | mRNA metabolism |
| POLR3C | RNA Polymerase III Subunit C | subunit of the RNA polymerase III complex | mRNA metabolism |
| PPM1F | Protein Phosphatase, Mg2+/Mn2+ Dependent 1F | Ser/Thr protein phosphatase |  |
| RBFOX1 | RNA Binding Protein, Fox-1 Homolog 1 | regulates tissue-specific alternative splicing |  |
| SEPT5 | Septin 5 | regulation of cytoskeletal organization and SNARE complexes | cytoskeleton |
| TBL1XR1 | Transducin Beta Like 1 X-Linked Receptor 1 | F-box-like protein involved in the recruitment of the ubiquitin/19S proteasome complex to nuclear receptor-regulated transcription units. | transcription |
| TTC27 | Tetratricopeptide Repeat Domain 27 | unknown |  |
| TUBA3C | Tubulin Alpha 3c | major component of microtubules | cytoskeleton |
| TUBGCP2 | Tubulin Gamma Complex Associated Protein 2 | microtubule nucleation at the centrosome | cytoskeleton |
| WASL | Wiskott-Aldrich Syndrome Like | Regulates actin polymerization | cytoskeleton |
| WWOX | WW Domain Containing Oxidoreductase | WW Domain Containing Oxidoreductase |  |

## Discussion

Here we present an integrated analysis of ADHD-associated CNVs identified in the first 11 published studies. The limited power of the individual studies has been a crucial bottleneck in the definition of high priority ADHD candidate genes for further studies. From the 1532 CNVs described in the 11 studies, using strict criteria, we extracted 2241 mRNA-coding genes; this number is likely to contain many false positives due to the individually low occurrence of the rare CNVs (20), but the number is too large to allow each CNV to be studied in (animal) models for validation and mechanistic insights. Here, we aimed to prioritize genes linked to ADHD among the 2241 based on robustness of findings across different bioinformatics approaches used in an integrative manner and including both human and animal model-derived data. For this, we selected only those genes that were recurrently affected by CNVs in the patients and focussed on the minimal overlapping region of different CNVs in a region. Furthermore, we removed all genes that were affected by a CNV in healthy controls of other studies, and those that were only partially duplicated (lacking a full coding sequence and promoter). Those stringent criteria substantially reduced the number of candidate genes from the CNVs to one fifth of the original number. We showed that the selected 432 high-priority genes were significantly more co-expressed in the developing brain in comparison to random gene groups; this is evidence that our selection enriches for biologically coherent genes that are expressed at the same time in the same tissue, a prerequisite for them to be involved in the same biological processes and cause similar phenotypes when disturbed.

Studies in model organisms can provide a wealth of phenotypic information, due to the high level of functional conservation across species. For ADHD, several model organisms have been shown to provide valid phenotypes. Those include monkey, rat, mouse, zebrafish, and fruit fly, where relevant phenotypes can be observed upon genetic manipulation or drug administration.(7) To identify biological processes underlying the selection of genes, we therefore took a novel approach that has not yet been applied to the field of neuropsychiatric disorders. We mined the Monarch Initiative database, which integrates genotype–phenotype relations across species, and found that 18 of the 432 high-priority genes, when manipulated in animal models, cause phenotypes that are face-valid to the ADHD core phenotypes attention-deficit, hyperactivity, and impulsivity. It is highly likely that the identified 18 genes whose protein-products form functional biological connections with genes/proteins for which detailed functional characterization, in particular information on ADHD-related phenotypes, is still lacking. We therefore retrieved direct interactors with the products of the 18 genes and identified an interconnected network of 66 proteins linked directly or indirectly to disease-relevant phenotypes (Figure 2). We also tested, whether our high-priority gene list itself formed protein-interaction modules with significantly interconnected proteins, which could provide us with information on biological processes. Indeed, we identified four modules comprising significantly connected proteins from our selection. Of these, the major modules, module 1 and module 4, connected to and were supported by ADHD-related phenotypes across species, showing the added value of cross-species analysis (Figure 2).

Module 1 contained two significantly linked proteins: WW-domain containing oxidoreductase (WWOX) and DNA-directed RNA polymerase I subunit RPA1 (POLR1A). WWOX, involved in autosomal recessive cerebellar ataxia-epilepsy-intellectual disability syndrome,(37) but no direct connections with ADHD are known. However, WWOX interacts with Protein phosphatase 1F (PPM1F), a Ser/Thr protein phosphatase that modulates RhoA and Ca^2+^/calmodulin-dependent protein kinase II pathways.(38,39) Other members of this phosphatase protein family are involved in mediating dopaminergic signalling via G-protein-coupled receptors.(40) In addition, miss-expression of PPM1F in the substantia nigra in Parkinson’s patients may implicate PPM1F more directly in dopaminergic biology.(41) Dopamine signalling pathways have been found altered repeatedly in ADHD patients and form the basis for the most widely used pharmacological treatment approach.(42,43) As we can directly connect PPM1F to the cross-species term hyperactivity and indirectly link *WWOX, POLR1A, POLR2B*, and *POLR3C* to ADHD core phenotypes, we can extend the network with genes potentially regulating dopaminergic signalling (Figure 2). The RNA polymerase II subunits (POLR1A, POLR2B, and *POLR3C*) are involved in the regulation and finetuning of transcription.

Module 2 clusters proteins required for blood–brain barrier formation, which function in cell–cell junctions and communication. This module contains one significantly connected protein: Catenin alpha-3 (CTNNA3). This adherence junction protein, also known to be associated with ASD, likely modulates cerebral and ependymal regions through GABA-A receptor activation.(44) Tight junction protein ZO-1 (TJP1) forms the connection to the other three proteins in this module. This gene is affected by CNVs in 28 ADHD cases, being the most frequently occurring gene affected by copy number alterations in our survey of the ADHD CNV studies (**Supplementary Table 1**). TJP1, together with Claudin-5 (CLDN5, affected in 12 ADHD cases) represents an important constituent of the blood–brain barrier.(45,46) Protein kinase C eta type (PRKCH) regulates TJP1.(47) Genes regulating neuronal cell adhesion are also significantly associated with ASD, schizophrenia, and bipolar disorder, raising the hypothesis that this mechanism plays a role across different neuropsychiatric disorders.(48,49) Due to the high number of ADHD cases with a CNV in this module, we postulate that cell–cell junctions play an important role in ADHD.

Module 3 contains two proteins that directly interact with each other: the significantly interconnected Density-regulated protein (DENR) and the Eukaryotic elongation factor 2 kinase (EEF2K), both involved in regulation and initiation of translation.(50) Regulation and initiation of translation has been linked to neuropsychiatric disorders including ADHD, through the regulation of brain-derived neurotrophic factor (BDNF).(51–53) Based on the repeated association and the described functional work, we suggest the involvement of transcriptional regulation as one of the mechanisms that modulate the ADHD risk.

Module 4 contains four significantly connected proteins: BIRC6, RAB15, SEPT5, and PHKB. Baculoviral IAP repeat-containing protein 6 (BIRC6 or Bruce) is an inhibitor of apoptosis involved in prostate cancer progression, but also acts in neuronal protection against apoptosis.(54,55) Ras-related protein Rab-15 (RAB15) is a direct connector of BIRC6, it plays a role in regulating synaptic vesicle membrane flow in nerve terminals.(56,57) Septin-5 (SEPT5) is involved in the binding of SNARE complexes, inhibiting synaptic vesicle exocytosis.(58) Recent studies have shown that manipulation of this gene in mice leads to altered social interaction and altered affective behaviours.(59,60) Phosphorylase b kinase regulatory subunit beta (PHKB) is involved in glycogen metabolism and has been linked to neuronal plasticity.(61) Other proteins in the hub link to the cross-species phenotype term hyperactivity and attention-deficit/hyperactivity disorder: MAPK1 is directly connected, and SEPT5, PARK2, MYC, and TUBA3C are indirectly connected. They are thus prime candidates for further evaluation in functional assays.

While this study started from rare CNVs, we found corroborating evidence for several of the genes implicated in ADHD also in studies of common genetic variants. Among the 26 CNV-affected genes most consistently observed across the different bioinformatics approaches applied in this study, we also found *RBFOX1* and *POLR3C* to be associated with the disorder in the largest SNP-based GWAS meta-analysis to date. In a recent study of Lee et al., *RBFOX1* was discovered to be the second most pleiotropic locus of the genome-wide meta-analysis amongst eight psychiatric disorders.(62) *RBFOX1* encodes a splice regulator which is regulating several genes involved in neuronal development and which is mainly expressed in the brain.(62–65) Animal models have shown that *RBFOX1* is involved in mouse corticogenesis and aggressive behaviours.(62,64,66,67) *POLR3C* is included in a well-known small CNV located at 1q21.1, which contributes to a broad spectrum of phenotypes in addition to ADHD, including morphological features and autism spectrum disorders (ASD).(68,69) Genes listed in Table 1 may be viewed as having, individually, the highest credibility as ADHD candidate genes. We therefore recommend to prioritize these genes in future studies searching for rare (single nucleotide) variants in ADHD and for functional characterization of gene-disease pathways. Table 1 shows that also among these 26 genes, common biological themes are present, such as transcription (already highlighted by the module analyse), mitochondria biology, mRNA metabolism, and cytoskeleton.

Our study should be viewed in the light of some strengths and limitations. We show that filtering based on recurrent CNVs restricted to ADHD cases in conjunction with complementary bioinformatics methods bares great potential to prioritise ADHD candidate genes. For the first time, cross-species phenotypes were used to identify candidate genes linked to ADHD core phenotypes, pointing to high-priority candidate genes. Several of those form significantly connected protein networks characterized by shared functions. The selection of genes analysed in this study is extensive, but is by no means exhaustive. First, it is important to note that by only analysing recurrent CNVs and focussing on the minimal regions of overlap amongst CNVs in the same region, genes with relevance for ADHD may be overlooked; on the other hand, the stringent evidence-based filtering holds high potential to uncover the most ADHD-relevant biological pathways. In addition, neither the surveyed CNV studies nor our study consider effects of CNVs onto the surrounding genetic landscape. A duplicated CNV translocation, for example, can have an impact on the expression of genes in the chromosomal region at the site of insertion, which alone or together with the duplicated genes can contribute to ADHD phenotypes.(70) Lastly, the criteria used for selection of CNVs were not identical across the 11 source studies.

In conclusion, our study shows that stringent filtering in CNV studies in combination with a complementary battery of bioinformatics approaches can identify ADHD candidate genes at increased levels of credibility. We suggest that testing genes operating in the identified modules and those 26 consistently found in all approaches employed here in animal models like rat, mouse, zebrafish, and fruit fly,(7) for their ability to modulate behaviour will provide further insights into the mechanistic pathways and biology of ADHD.

---

**Supplementary Figure 1:** Regional association of genes most strongly associated with ADHD. Regional association plots showing association signal for ADHD in the PGC-iPSYCH GWAS meta-analysis data for the two most strongly associated high priority ADHD candidate genes, including flanking regions of 100kb. (A) *POLR3C* locus with the top-SNP (rs376814422) indicated by the black arrow. (B) *RBFOX1* locus with the top-SNP (rs6500945) indicated by the purple dot. Results are shown as -log (p-value) for genotyped and imputed SNPs. The colour of each marker reflects its LD (r2) with the SNP indicated by the purple dot. The recombination rate is plotted in blue. cm/Mb, centimorgan/megabase. Chr, chromosome.

**Supplementary Table 1:** Full lists of raw input data points extracted from 11 ADHD CNV studies (Tab 1), high-priority gene selection (Tab 2) and low-priority gene selection (Tab 3).

**Supplementary Table 2:** Full list of all selected ADHD core phenotype labels in the Monarch Initiative database query.

**Supplementary Table 3:** Overview of all included studies reporting rare CNVs in an ADHD cohort.

**Supplementary Table 4:** Significant connected genes of the high-priority gene list after DAPPLE analysis and the assigned modules in Figure 3.

**Supplementary Table 5:** Results MAGMA analysis of the 26 genes consistently observed in all the different approaches

**Supplementary Methods 1:** SQL query script for the UCSC database, GWAS meta-analyses data set for ADHD and Gene-based association analyses for ADHD GWAS meta-analyses data

## Author contribution

BH: performed data analysis and created interaction networks, contributed to manuscript writing. MV: conceived the study, performed data analysis, processed CNVs using SQL, curated cross-species Monarch Initiative and BioGRID information, contributed to supervision and manuscript writing. MK: Performed gene-based association analyses and helped with data interpretation. PC and MF: performed the co-expression analysis. BF: contributed to supervision and manuscript writing. AS: conceived the study, contributed to supervision and manuscript writing. All authors proof-read the manuscript.

## Acknowledgements

Benjamin Harich was funded by a Radboud University Medical Center PhD grant. Monique van der Voet’s research was supported by the Netherlands Organization for Scientific Research (NWO) VENI grant (grant 91.614.084). Marieke Klein was supported by a grant from the Netherlands Science Agenda (NWA) for the NeuroLabNL project (grant 400.17.602). Barbara Franke’s contribution was supported by a personal Vici grant from the Netherlands Organization for Scientific Research (NWO; grant 016-130-669). This work also received support from the European Community’s Horizon 2020 (H2020/2014 – 2020) European Trainings Network Programme MiND under grant agreement n° 643051 to Barbara Franke and Annette Schenck. Additional support was received from the ECNP Network ADHD across the Lifespan and the European Community’s Horizon 2020 Programme under grant agreements n° 667302 (CoCA) and n° 728018 (Eat2beNICE). This report reflects only the authors’ views, and the European Union is not liable for any use that may be made of the information contained therein.

