## Supplementary Table 1 for "From rare Copy Number Variations to biological processes in ADHD"

**Supplementary Table 3 - Cohort Studies**

| Study | Cohort | Cases (n) | Controls (n) | CNV size |
|---|---|---|---|---|
| 2010 Elia | Caucasian (Philadelphia) | 335 trios | 2026 | incl. Small |
| 2010 Williams | Caucasian (UK) | 366 | 1047 | > 500kb |
| 2011 Lesch | Caucasian (Germany) | 99 + parents | 100 | one 75kb, the rest larger |
| 2011 Lionel | Caucasian (Ontario, Germany) | 248 | 2357 | >20kb |
| 2012 Elia | Caucasian (Philadelphia) | 1013 | 4105 | incl. Small |
| 2012 Stergiakouli | Caucasian (Wales, Scotland, Ireland) | 727 | 5081 | > 500kb |
| 2012 Wiliams | Caucasian (European) | 732 | 2455 | > 100kb |
| 2014 Jarick | Caucasian (Germany) | 489 | 1285 | > 500kb |
| 2014 Martin | Caucasian (Wales, Scotland, Ireland) | 727 | 5081 | > 500kb |
| 2014 Ramos-Quiroga | Caucasian (Spain) | 400 | 526 | > 100kb (except Table S4) |
| 2013 Yang | Asian (China) | 1040 | 963 | > 100kb |
