## Supplementary Table 3 for "From rare Copy Number Variations to biological processes in ADHD"

**Supplementary Table 2 - Monarch Initiative Labels**

| Search term in Monarch Initiative database |
|---|
| hyperactive |
| Impulsivity |
| attention |
| hyperactive |
| attention |
| Impulsivity |
| increased vertical activity |
| Hyperactivity |
| Hyperactivity |
| behavioral |
| Impulse |
| locomot |
| hyperactive |
| hyperactive |
| locomot |
| increased frequency |
| increased rate |
| behavioural |
| increased occurrence |
| locomotory behavior |
| increased speed locomotion |
| increased occurrence |
| increased rate |
| increased rate |
| locomot |
| behavioural |
| increased occurrence |
| increased occurrence |
| increased frequency |
| increased frequency |

**Results**

| object | object_label | Monarch genes 12-jan-2017 |
|---|---|---|
| FBcv:0000392 | hyperactive | 47 |
| HP:0000734 | Disinhibition | 36 |
| HP:0000736 | Short attention span | 125 |
| HP:0000752 | Hyperactivity | 684 |
| HP:0007018 | Attention deficit hyperactivity disorder | 114 |
| HP:0100710 | Impulsivity/abnormal impulsive behavior control | 62 |
| MP:0002574 | increased vertical activity | 77 |
| MP:0002629 | hyperactivity elicited by ethanol administration | 8 |
| MP:0008911 | induced hyperactivity | 43 |
| MP:0009751 | enhanced behavioral response to alcohol | 17 |
