## Supplementary Table 4 for "From rare Copy Number Variations to biological processes in ADHD"

**Supplementary Table 4 - DAPPLE Module Significance**

| Gene | P corrected | Assigned Module in Figure 3 |
|---|---|---|
| WWOX | 0,0297 | Module 1 |
| POLR1A | 0,0356 | Module 1 |
| CTNNA3 | 0,0415 | Module 2 |
| DENR | 0,0317 | Module 3 |
| BIRC6 | 0,0120 | Module 4 |
| RAB15 | 0,0396 | Module 4 |
| SEPT5 | 0,0396 | Module 4 |
| PHKB | 0,0139 | Module 4 |
| SDK1 | 0,0020 | no assigned module |
| UXS1 | 0,0020 | no assigned module |
| C12orf65 | 0,0020 | no assigned module |
| PRRT2 | 0,0080 | no assigned module |
| FAM84B | 0,0100 | no assigned module |
| TAGLN2 | 0,0159 | no assigned module |
| FIS1 | 0,0179 | no assigned module |
| SLC6A12 | 0,0219 | no assigned module |
| ADSS | 0,0278 | no assigned module |
| ATP1A4 | 0,0317 | no assigned module |
| FKBP3 | 0,0337 | no assigned module |
| SETD8 | 0,0454 | no assigned module |
