## Supplementary Text 1 for "From rare Copy Number Variations to biological processes in ADHD"

### SQL query

```
#select below for SQL query:

select
CNV.name as Study,
CNV.type as Type,
Gene.name as Refseq,
Gene.name2 as Name,
Gene.strand as Strand,
CNV.chrom as Chromosome,
CNV.chromStart as CNV_Start,
CNV.chromEnd as CNV_End,
Gene.txStart as Gene_Start,
Gene.txEnd as Gene_End,
(Gene.txEnd-Gene.txStart) as Gene_Size,
(CNV.chromEnd-CNV.chromStart) as CNV_Size,

#CREATE COLUMN LOCATION
if  (CNV.chromStart<Gene.txEnd and Gene.txStart<CNV.chromEnd,
#The CNV overlaps with the gene, where does it overlap?
if  (Gene.txStart<CNV.chromStart,
#It overlaps with right, what part of gene?
if (Gene.strand='+',"Cterm","Nterm"),
if (CNV.chromEnd<Gene.txEnd,
#It overlaps with left, what part of gene?
if (Gene.strand='+',"Nterm","Cterm"),
#Neither left or right, thus whole gene
"CDS")
),
#The CNV does not overlap with the gene, does it overlap with 2kb of regulatory region:
if(Gene.txEnd<CNV.chromStart,if(Gene.strand='+',"3'UTR","PROM"),if(CNV.chromEnd<Gene.txSt
art,if(Gene.strand='+',"PROM","3'UTR"),"NULL"))


) as CNV_Location,

#CREATE COLUMN OVERLAP
if (CNV.chromStart<Gene.txEnd and Gene.txStart<CNV.chromEnd,
#The CNV overlaps with the gene, how large is the overlap?
if (Gene.txStart<CNV.chromStart,Gene.txEnd-CNV.chromStart,
if (CNV.chromEnd<Gene.txEnd,CNV.chromEnd-Gene.txStart,Gene.txEnd-Gene.txStart)
),
#The CNV does not overlap with the gene:
"0") as Coding_Overlap,

#CREATE COLUMN GENEPERCENTAGE
```

```
if (CNV.chromStart<Gene.txEnd and Gene.txStart<CNV.chromEnd,
#The CNV overlaps with the gene, how large is the overlap?
if (Gene.txStart<CNV.chromStart,(Gene.txEnd-CNV.chromStart)/(Gene.txEnd-Gene.txStart),
if (CNV.chromEnd<Gene.txEnd,(CNV.chromEnd-Gene.txStart)/(Gene.txEnd-Gene.txStart),"1")
),
#The CNV does not overlap with the gene:
"0") as Coding_Fraction,

#CREATE COLUMN PROMOTER
if(Gene.strand='+',if(CNV.chromStart<Gene.txStart,Gene.txStart-
CNV.chromStart,"NULL"),if(Gene.txEnd<CNV.chromEnd,CNV.chromEnd-Gene.txEnd,"NULL"))
as "5'UTR_Overlap",

#CREATE COLUMN UTR
if(Gene.strand='-',if(CNV.chromStart<Gene.txStart,Gene.txStart-
CNV.chromStart,"NULL"),if(Gene.txEnd<CNV.chromEnd,CNV.chromEnd-Gene.txEnd,"NULL"))
as  "3'UTR_Overlap",

#CREATE COLUMN GAP
if(cnv.chromend<gene.txstart,gene.txstart-
cnv.chromend,if(gene.txend<cnv.chromstart,cnv.chromstart-gene.txend,"NULL"))
as  "Gap_CNV-Gene"

# DECLARE THE TABLE:
##########################################################################

from 140822_transcoded_corrected

#####################################################################################
############
as CNV

left join refGene as Gene on

(CNV.chrom=Gene.chrom and not(Gene.txEnd<(CNV.chromStart-2000) or
(CNV.chromEnd+2000)<Gene.txStart))
```

**GWAS meta-analyses data set for ADHD**

The cohorts include eleven clinical collections of the PGC and 37,076 samples of the Danish Bloodspot efforts (iPSYCH). Samples were of Caucasian or Han Chinese origin and met diagnostic criteria according to the DSM-IV.[1] Written informed consent was obtained from all participants. Each study was approved by the respective institutional review board or local ethics committee. The meta-analytic data used in this study were available as summary statistics, including genome-wide SNP data with corresponding P-values and odds ratios. Detailed procedures of DNA isolation, whole-genome genotyping and imputation were described previously.[1] Shortly, genome-wide data was obtained from different genotyping arrays and was imputed using 1000 Genomes data as a reference panel (phase 3, version 5 (1KGP3v5)) in NCBI build 37 (hg19) coordinates) for autosomal SNPs.[2] Meta-analytic data were processed through a stringent quality control pipeline applied at the PGC.[1] Only SNPs with an imputation quality score of INFO $\geq 0.8$ and a minor allele frequency $\geq 0.01$ were included in our analyses.

**Gene-based association analyses for ADHD GWAS meta-analyses data**

We used data from the recent meta-analysis of genome-wide association studies (GWAS) of 20,183 patients with ADHD and 35,191 controls as performed by the Psychiatric Genomics Consortium (PGC) ADHD Working Group and the Danish iPSYCH Initiative. Details on the samples and quality control (as described above and in Demontis et.al 2019).[1]

Gene-based association analyses were performed using the Multi-marker Analysis of GenoMic Annotation (MAGMA) software package (version 1.05)[3]. First, genome-wide SNP data from a reference panel (1000 Genomes, v3 phase1)[4] was annotated to NCBI Build 37.3 gene locations using a symmetric 100 kb flanking window and both files were downloaded from http://ctglab.nl/software/magma. Next, the gene annotation file was used to map the genome-wide SNP data, to assign SNPs to genes, and to calculate gene-based p-values. For the gene-based analyses, single SNP p-values within a gene were transformed into a gene-statistic by taking the mean of the χ2-statistic among the SNPs in each gene. To account for linkage disequilibrium (LD), the 1000 Genomes Project European sample was used as a reference to estimate the LD between SNPs within (the vicinity of) the genes (http://ctglab.nl/software/MAGMA/ref_data/g1000_ceu.zip). Gene-wide p-values were converted to z-values reflecting the strength of the association of each gene with the phenotype, with higher z-values corresponding to stronger associations. All individual genes were investigated, by reviewing their gene test-statistics. Genes were considered gene-wide significant if they reached the Bonferroni correction threshold-adjusted for the number of genes tested ($p<0.05/26$).

1. Demontis, D. *et al.* Discovery of the first genome-wide significant risk loci for attention deficit/hyperactivity disorder. *Nat. Genet.* **51**, 63–75 (2019).
2. Auton, A. *et al.* A global reference for human genetic variation. *Nature* **526**, 68–74 (2015).
3. de Leeuw, C. A., Mooij, J. M., Heskes, T. & Posthuma, D. MAGMA: Generalized Gene-Set Analysis of GWAS Data. *PLoS Comput. Biol.* **11**, 1–19 (2015).
4. Consortium, T. 1000 G. P. A map of human genome variation from population scale sequencing. *Nature* **467**, 1061–1073 (2010).
