## Supplementary figures and images for "From rare Copy Number Variations to biological processes in ADHD"

### Supplementary Figure 1

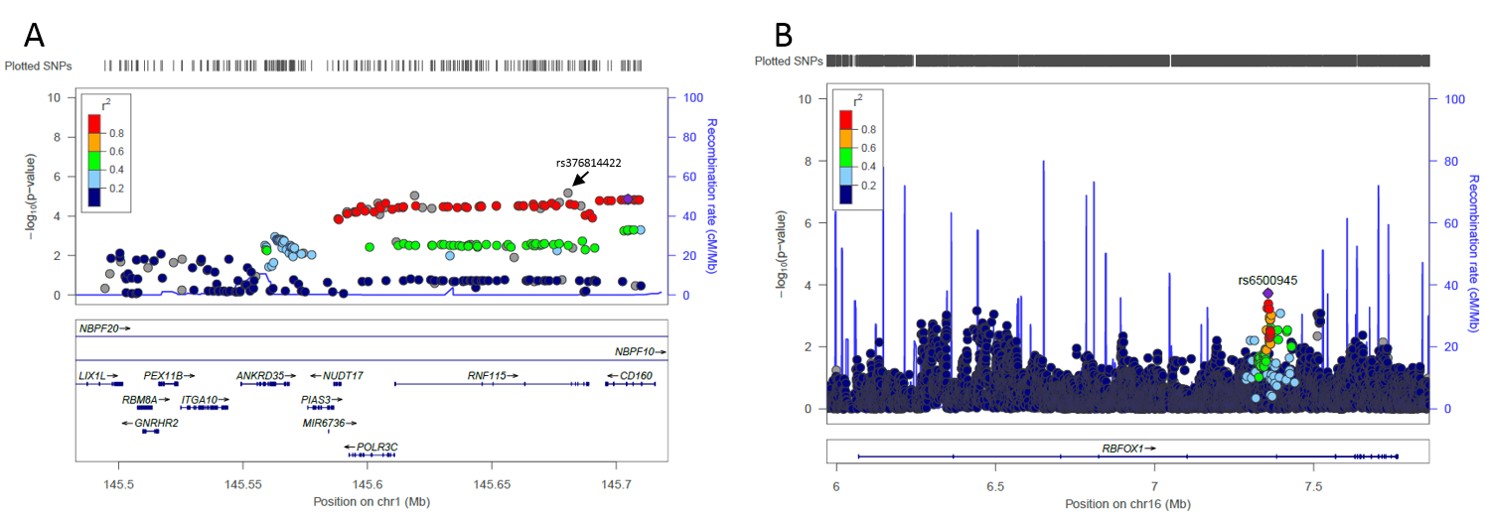
